# Coral skeletal cores as windows into past Symbiodiniaceae community dynamics

**DOI:** 10.1101/2025.03.21.644536

**Authors:** Jose F. Grillo, Vanessa Tirpitz, Jessica Reichert, Marine Canesi, Stéphanie Reynaud, Eric Douville, Maren Ziegler

## Abstract

The symbiosis between the dinoflagellate Symbiodiniaceae family and reef-building corals underpins the productivity of coral reefs. This relationship facilitates the deposition of calcium-carbonate skeletons that build the reef structure thanks to the energy derived from photosynthesis. The loss of Symbiodiniaceae from coral tissues—resulting in coral bleaching—impedes coral growth and can lead to mass mortality if the symbiosis fails to recover. Given that Symbiodiniaceae communities are dynamic and can shift in response to environmental stressors in the decades- to centuries-long lifespan of coral colonies, understanding these changes is crucial. Although the reconstruction of Symbiodiniaceae communities from coral skeleton records has recently been demonstrated as feasible, no studies have yet assessed reconstructions across different species and locations. Here, we present an approach to use coral skeletons for reconstructing the Symbiodiniaceae community on decadal and centennial scales and resolving dynamics related with coral species and environmental history of sampling locations. For this, we used dated coral skeleton cores from *Porites Iobata* and *Diploastrea heliopora*, species commonly used as climate archives, sampled in Palau and Papua New Guinea. We also examined the effect of various DNA extraction protocols on community reconstruction. Here we show that the reconstructed Symbiodiniaceae communities significantly varied across all cores and DNA extraction methods, with decalcification-based protocols enhancing the retrieval of skeletal-bound DNA. Moreover, we observed distinct community dynamics related to the specific coral host and sampling location. Notably, associations of Symbiodiniaceae dynamics with past heat stress events were apparent in cores of both species from Palau. Our findings enable a deeper understanding of the temporal and spatial variability in Symbiodiniaceae communities, offering insights that may refine the use of paleobiological proxies in climate studies and reveal broader ecological trends and microbially-aided adaptation pathways in corals.

## 1. Introduction

The health and productivity of coral reefs builds on the symbiosis of reef-building corals with photosynthetic dinoflagellates from the diverse Symbiodiniaceae family that reside within their tissues (Nitschke et al. 2022; Helgoe et al. 2024). The translocation of carbohydrates produced by Symbiodiniaceae during photosynthesis is used by the coral to deposit their characteristic calcium carbonate (CaCO3) skeleton, which builds the reef framework (Al-Horani 2015). At the same time, this symbiosis between corals and Symbiodiniaceae is also their Achilles’ heel due to its susceptibility to heat stress (Helgoe et al. 2024). Increasing duration, frequency, and severity of marine heat waves cause coral bleaching which is the loss of the Symbiodiniaceae from coral tissue (Frölicher et al. 2018; Helgoe et al. 2024). Coral bleaching results in starvation, reduced skeletal growth, and may lead to mortality (Helgoe et al. 2024). Marine heatwaves promoted by climate change have caused mass bleaching events which led to the loss of between 30 and up to 90 % coral cover in reef ecosystems across the Pacific Ocean and the Arabian Seas (Riegl et al. 2018; Raymundo et al. 2019). More recently, the Fourth Global Coral Bleaching Event was recorded in 2024 and while its impact is still being studied, future global-scale bleaching events are expected to continue with rising CO2 emissions (Reimer et al. 2024).

Symbiodiniaceae are a diverse family, whose members are functionally distinct regarding environmental niche space and the traits they confer to the coral (LaJeunesse et al. 2018). These functions are assumed to be determined by one or few abundant symbiont taxa that dominate the community along with several background symbionts that usually persist at low abundance (Quigley et al. 2014; Davies et al. 2023). Symbiodiniaceae species differ in the amount of carbon they translocate to the coral, influencing their host’s growth rates (Pettay et al. 2015), and they notoriously differ in heat stress tolerance (Swain et al. 2017; Xiao et al. 2022). So-called adaptive bleaching (Buddemeier and Fautin 1993), a dynamic process in which the main symbiont taxon is replaced, may rescue the coral after bleaching and may subsequently result in higher heat stress tolerance (Baker et al. 2004; Boulotte et al. 2016). The bleached symbionts may be replaced from the extant community through shuffling or recruited from the environment through switching (Baker 2003; Quigley et al. 2022). As such, the study of Symbiodiniaceae through tissue or mucus sampling provides a snapshot of the community at particular time points that are limited by availability of samples (Million et al. 2024). Extraction of historical DNA from dried museum specimens can extend surveys back in time (Rowe et al. 2011). A study conducted on up to 172 year old soft corals in museum collections suggested that Symbiodiniaceae communities remained largely unchanged compared to recent samples (Baker et al. 2013). Yet, approaches that rely on tissue samples fail to reveal long-term dynamics following periods of environmental stress throughout the century-scale lifespan of reef-building corals.

Coral skeletons have been extensively used for the reconstruction of past environmental conditions through analyses of stable isotopes, trace elements, and biogeochemistry of the skeletal material (Thompson 2022; Canesi et al. 2024). The signals recorded in coral skeletons over centuries are proxies for past climates, ocean chemistry, and pollution (Ramos et al. 2009; Thompson 2022; Canesi et al. 2024). Recently, ancient DNA (aDNA) applications probed sub-fossil deep sea coral skeletons for species identification (Waller et al. 2007) and subsequently to identify genomic similarities between millenia-old and living colonies of *Acropora palmata* and their bacterial communities (Scott et al. 2022). The skeletons of tropical stony corals have successfully been used for reconstructions of host phylogenies and of prokaryotic and Symbiodiniaceae communities (Scott et al. 2022; González-Pech et al. 2024; Rey et al. 2025). Nevertheless, the use of single cores is associated with inherent limitations that has prevented the study of Symbiodiniaceae community changes related to geographic and coral host-associated sources of variability that remain to be explored in future studies.

The overarching goal of this study was to evaluate the use of skeleton cores from massive coral species for the reconstruction of Symbiodiniaceae communities at decadal and centennial time scales across species and locations. For this, century-old coral skeleton cores were sampled from two massive species (*Porites lobata* and *Diploastrea heliopora*) from two locations in the Pacific (Papua New Guinea and Palau) and ancient Symbiodiniaceae community dynamics were analyzed using ITS2 amplicon sequencing. Our specific aims were to: 1) assess the effect of three different DNA extraction methods on the reconstructed Symbiodiniaceae communities, 2) evaluate changes in reconstructed communities obtained by species and sampling location, and 3) probe differences in the reconstruction dynamics between two massive growing coral species in the context of past potential heat stress events. The results generated by this study highlight the necessity of moving beyond proof-of-concept studies to explore region and coral-specific community dynamics at broader scales needed to fine-tune survivorship projections and coral restoration efforts based on long-term time series data derived from coral skeletons.

## 2. Materials and Methods

### 2.1. Collection of coral cores

Four coral cores were drilled from colonies located in North Palau (7°47’ N, 134°35’ E) and Losuia Tabungora Island in Papua New Guinea (9°21’ S, 152°02’ E) during the Tara Pacific expedition (Planes et al. 2019) in 2017 and 2018, respectively (Fig. 1). The cores were collected as described in Canesi et al. (2023, 2024) for geochemical reconstruction of past environmental conditions and stored at room temperature after sampling. Briefly, a 40-150 cm long skeletal core was taken with a hydraulic drill (Stanley Black & Decker, US) with a 7 cm diameter corer from one living colony of *Porites lobata* (Dana, 1846) and *Diploastrea heliopora* (Lamark, 1816) per site at 13-16 m. The cores were rinsed with fresh water at the time of collection, left to air dry, and stored at room temperature until processing. *Porites lobata* is one of the species most commonly used for reconstruction of climate pasts and has a relatively high linear extension rate of 7.5 to 16.3 mm per year (Klein and Loya 1991; Edinger et al. 2000). *Diploastrea heliopora* is a comparably slow growing species (ca. 2-4 mm per year) which enables long-term time series reconstruction (Masuda et al. 2010; Canesi et al. 2024).

**Figure 1.**
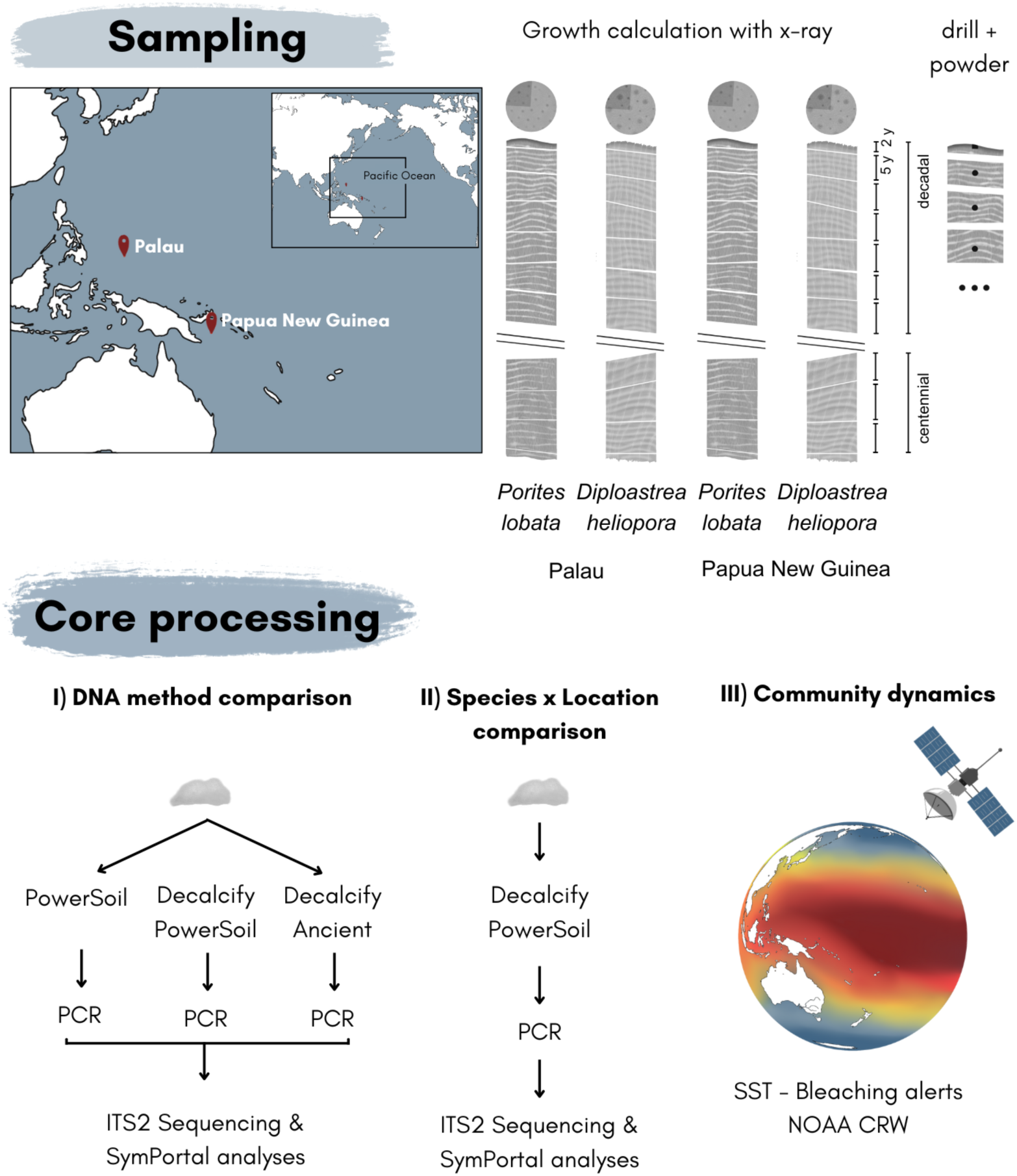
Overview of study design. Coral cores of the species *P. lobata* and *D. heliopora* were sampled in Palau and Papua New Guinea. Fragments along the growth axis were drilled to I) compare DNA extraction methods, II) compare symbiont communities across species and sampling locations, and III) link community dynamics to bleaching alerts of the past century.

### 2.2. Experimental design

To test the effect of different DNA extraction methods on the reconstruction of Symbiodiniaceae communities from coral skeleton cores three methods were compared in this study based on commercially available kits and the presence of decalcification steps prior to the DNA isolation (Fig 1). Moreover, two time frames were chosen to reconstruct the Symbiodiniaceae community dynamics. The first 30 years before sampling were analyzed to study the recent dynamics of Symbiodiniaceae communities after potential bleaching events determined by combining sea surface temperature records and bleaching alerts at each of the sampling locations by the NOAA Coral Reef Watch dating back to 1985 (Skirving et al. 2020). Moreover, 80 to 95 year-old fragments of *P. lobata* and 100 to 115 year-old fragments of *D. heliopora* were sampled to assess the effectiveness of century-scale Symbiodiniaceae community reconstructions.

### 2.3. Coral core sample processing

Skeletal cores were halved along the longitudinal axis, dated through X-ray and the top of each core containing the tissue layer was sectioned including the skeletal fraction corresponding to two years. All following older sections were cut into 5-year fragments based on a calculation of linear extension rates, where one year was assigned as the combination of a high and low density band, and one half was further halved to obtain a fragment representing one longitudinal quarter of the original 7 cm-diameter cylindrical core. All cores were treated with UV light for 15 mins for surface decontamination prior to sample processing. The coring equipment (drill bit, ceramic mortar, and pestle) and all working surfaces were sterilized. Approximately 1 g of skeleton was taken from each 5-year fragment with a sterile metallic drill bit (diameter: 6 mm) and pulverized with a sterile ceramic mortar and pestle. Ten samples per coral core were used, resulting in a total of 40 skeletal samples from the two coral species from two locations.

### 2.4. Molecular sample processing

A comparison of DNA extraction methods was performed with the 10 powder samples of the *P. lobata* core from Papua New Guinea to test the effect of the extraction method on the reconstruction of Symbiodiniaceae communities in ancient coral samples. These tests included an extraction blank for each processing batch. Three different types of extraction were compared: 1) DNA was extracted from a 250 mg stock sample using the PowerSoil Pro kit (Qiagen, Germany) following the manufacturer’s instructions with a final elution volume of 50 µL. Methods 2 and 3 shared a decalcification step, which was not part of method 1. 2) A 250 mg stock sample was taken, first decalcified, and then the DNA was extracted with the same PowerSoil Pro kit (Qiagen, Germany). For decalcification, sterile-filtered EDTA solution 0.5 M (pH 7.5) was added to the skeletal powder and incubated at 37 °C on a Thermomixer comfort (Eppendorf, Germany) at 500 RPM until all skeletal material was dissolved. 3) The ancient DNA extraction protocol proposed by (Rohland et al. 2018) based on a decalcification step, the use of commercial silica spin columns and custom buffers that allow the recovery of short DNA fragments (as small as 35 bp) was tested with a 50 mg sample and the binding buffer D. Briefly, the coral skeleton powder is decalcified and lysed with a buffer containing EDTA 0.5M (pH 8.0), 10 mg/mL proteinase K and Tween 20. The lysate supernatant is added to the binding buffer D (5 M guanidine hydrochloride, 40% (vol/vol) 2-propanol, 0.12 M sodium acetate and 0.05% (vol/vol) Tween 20) and transferred to the silica spin columns. The spin columns are washed with PE buffer (Qiagen, Germany) and the DNA is eluted with a custom-made buffer based on 1 M Tris-HCl (pH 8.0), EDTA 0.5M (pH 8.0) and Tween 20.

The samples from the remaining three cores were extracted using only the decalcification and Powersoil Pro approach following method 2) above to examine differences in the Symbiodiniaceae community dynamics between coral species and sampling locations. For the PCRs, 6.4 µL of DNA template was used with the Symbiodiniaceae-specific primers SYM_VAR_5.8S2 and SYM_VAR_REV (Hume et al. 2018) and the Multiplex PCR Master Mix (Qiagen, Germany) with a final volume of 40 µL. One negative control per PCR batch was included during the sample processing. PCRs were cleaned using the QIAquick PCR purification kit (Qiagen, Germany). The cleaned PCR products were sent for library preparation and sequencing on the DNBseq G400 sequencing platform (BGI, Hong Kong).

### 2.5. Sequence data processing

Demultiplexed, quality-controlled reads were submitted to the SymPortal analytical workflow (Hume et al. 2019). SymPortal was designed for the analysis of Symbiodiniaceae communities through the recognition of co-occurring intragenomic sequences within each sample as so-called type profiles (Hume et al. 2019). Given the expected DNA degradation in the ancient DNA samples used here, we did not assume that all sequences necessary to distinguish SymPortal type profiles would be reliably recovered in the samples. In addition, the unknown mechanism of DNA or symbiont entrapment in the coral skeleton violates the SymPortal assumption of one Symbiodiniaceae species per genus in a sample. Therefore, post-MED sequences were used for all analyses. All sequences were deposited under the NCBI bioproject ID XXX and accessible in the SymPortal Database at XXX.

### 2.6. Data analyses

All analyses, visualizations, and statistical tests were performed in the R statistical software (version 4.3.2, R Core Team 2023). The absolute abundance count table of the post-MED sequences generated by the SymPortal workflow was pre-processed by removing all Symbiodiniaceae sequences present in the negative control samples for the DNA extraction and PCR negative control corresponding to each processing batch. Because of the known susceptibility of aDNA approach to contamination (Yang and Watt 2005; Hofreiter et al. 2021), we chose this stringent approach. This resulted in an average of 39,298 post-MED sequences per sample (range: 16,940-110,234; Table S1). The relative abundance of post-MED sequences per sample was then calculated and only sequences with a relative abundance > 0.1 % over all samples were retained in the analysis. The relative abundance of Symbiodiniaceae sequences per sample were visualized with a stacked bar plot using tidyverse (version 2.0.0, Wickham et al. 2019) and the ggnested package for color palette generation (version 0.1.1). The Symbiodiniaceae communities of each sample were visualized with an ordination plot based on non-Metric Multidimensional Scaling (nMDS) of Bray-Curtis distances with *metaMDS* from the vegan package (version 2.6-8, Oksanen et al. 2024). Finally, the number of shared Symbiodiniaceae sequences were visualized with an UpSet plot with the ComplexUpset package (version 1.3.3, Krassowski et al. 2022). Separate Permutational Multivariate Analysis of Variance (PERMANOVA, Anderson 2017) based on Bray-Curtis distances were performed to test for differences between the ITS2 communities obtained with different DNA extraction methods for the *P. lobata* core from Papua New Guinea and for differences between coral species and sampling locations in a fully crossed design, respectively. Both statistical tests were computed with 999 permutations and multivariate homogeneity of variance was tested with the vegan package. Post hoc permutational pairwise comparisons were computed with the pairwise.adonis package (version 0.4.1, Martinez Arbizu 2017).

The daily resolution sea surface temperature (SST) and bleaching alert data for both sampling locations obtained from the NOAA CRW (accessed on 18-02-2025, (Skirving et al. 2020) was filtered to coincide with the 5 year intervals of the coral skeleton core and visualized as a bar plot. This metric distinguishes three bleaching alert levels, which correspond to risk of reef- wide bleaching (level 1), risk of reef-wide bleaching with mortality of heat-sensitive corals (level 2), and risk of multi-species mortality (level 3).

## 3. Results

### 3.1. Symbiodiniaceae communities differed according to the DNA extraction protocol

The reconstruction of Symbiodiniaceae communities from coral skeleton cores was successful for the three DNA extraction protocols at decadal and centennial time scales, although differences were observed between the resulting communities (PERMANOVA, Pseudo-F = 1.7595, p < 0.001, Fig. 2A,C-E, Table 1, Table S2). Of note, the two protocols including the decalcification step were more similar to each other than to the protocol lacking this step (Table S2). While they had similar multivariate homogeneity of variances (Betadisper, p > 0.05, Table S3), the overall patterns were also reflected in the high number of 421 to 435 unique post-MED sequences per DNA extraction method (Fig. 2B). The protocols with a decalcification step shared a higher number of sequences (67), than each of them with the third protocol (51 and 63) or all protocols (37) (Fig. 2B).

**Figure 2.**
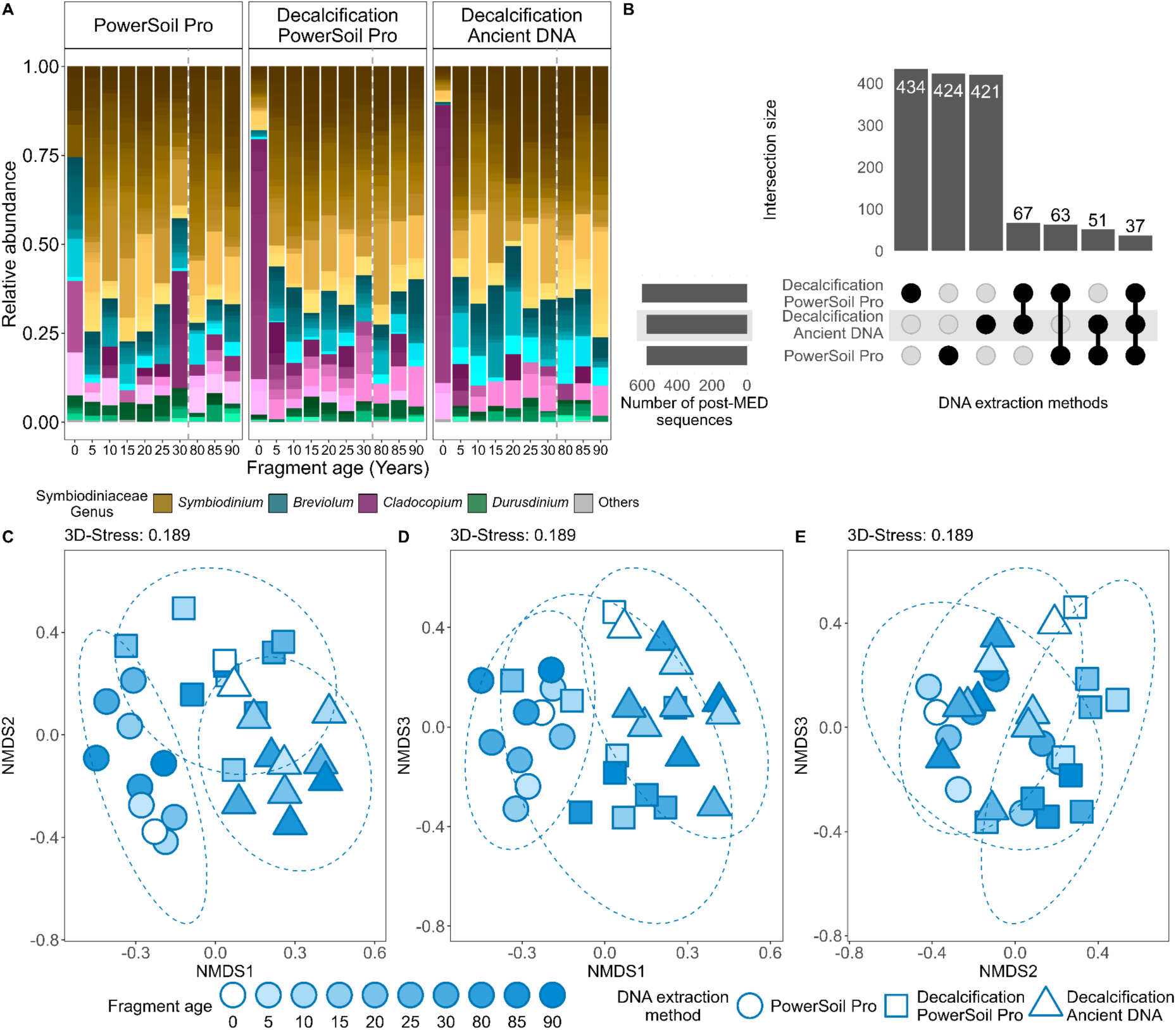
Symbiodiniaceae community from one *P. lobata* skeleton core collected from Papua New Guinea reconstructed with three different DNA extraction methods. **A**. Relative abundance of post-MED Symbiodiniaceae sequences per genus. The shades of color represent unique post-MED sequences. **B**. Number of shared post-MED sequences between the DNA extraction methods. The intersection size (vertical bar plot) represents the number of shared sequences of a set of DNA methods represented by filled points. **C-E**. Ordination of Symbiodiniaceae community composition with Non-metric Multidimensional Scaling based on Bray-Curtis distances of post–MED sequences. Ellipses represent 95 % CI.

**Table 1.**
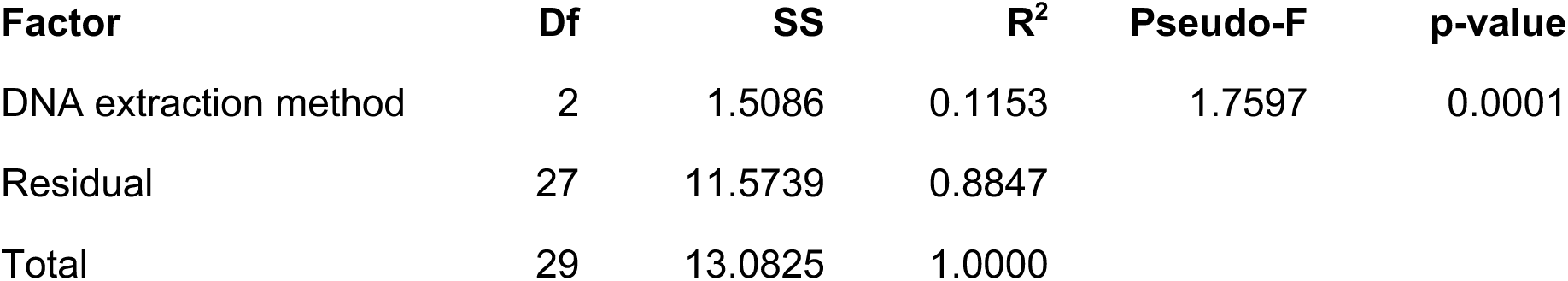
Statistical test of differences in the reconstructed Symbiodiniaceae communities obtained with three DNA extraction protocols from a *P. lobata* core sampled in Papua New Guinea. PERMANOVA on Bray-Curtis distances with 999 permutations. DF: degrees of freedom, SS: sum of squares, Pseudo-F: pseudo-F statistic.

The Symbiodiniaceae genera *Symbiodinium*, *Breviolum*, *Cladocopium,* and *Durusdinium* were recovered from all coral core samples and all DNA extraction methods, although community composition differed between protocols (Fig. 2A, C-E). Post-MED sequences in the genus *Symbiodinum* were most diverse and they also represented the most abundant fraction in all skeletal samples, but not in the top skeletal sample that included the tissue layer, where *Cladocopium* tended to be more abundant for the protocols employing a decalcification step (Fig. 2A).

### 3.2. Reconstructed Symbiodiniaceae community dynamics differed by species and location

Symbiodiniaceae communities differed between coral species (PERMANOVA, Pseudo-F = 1.63, p < 0.001), sampling locations (PERMANOVA, Pseudo-F = 1.69, p < 0.001), and between all cores across species and locations (PERMANOVA, Pseudo-F = 2.2304, p < 0.001, Fig. 3, Table S4, Table S5). Symbiodiniaceae communities between the *P. lobata* cores differed in multivariate dispersion (Betadisper, p < 0.05, Table S6), which was driven by the similarity of the core top fragments with tissue, while the older skeletal samples had distinct communities between cores (Fig. 3D).

**Figure 3.**
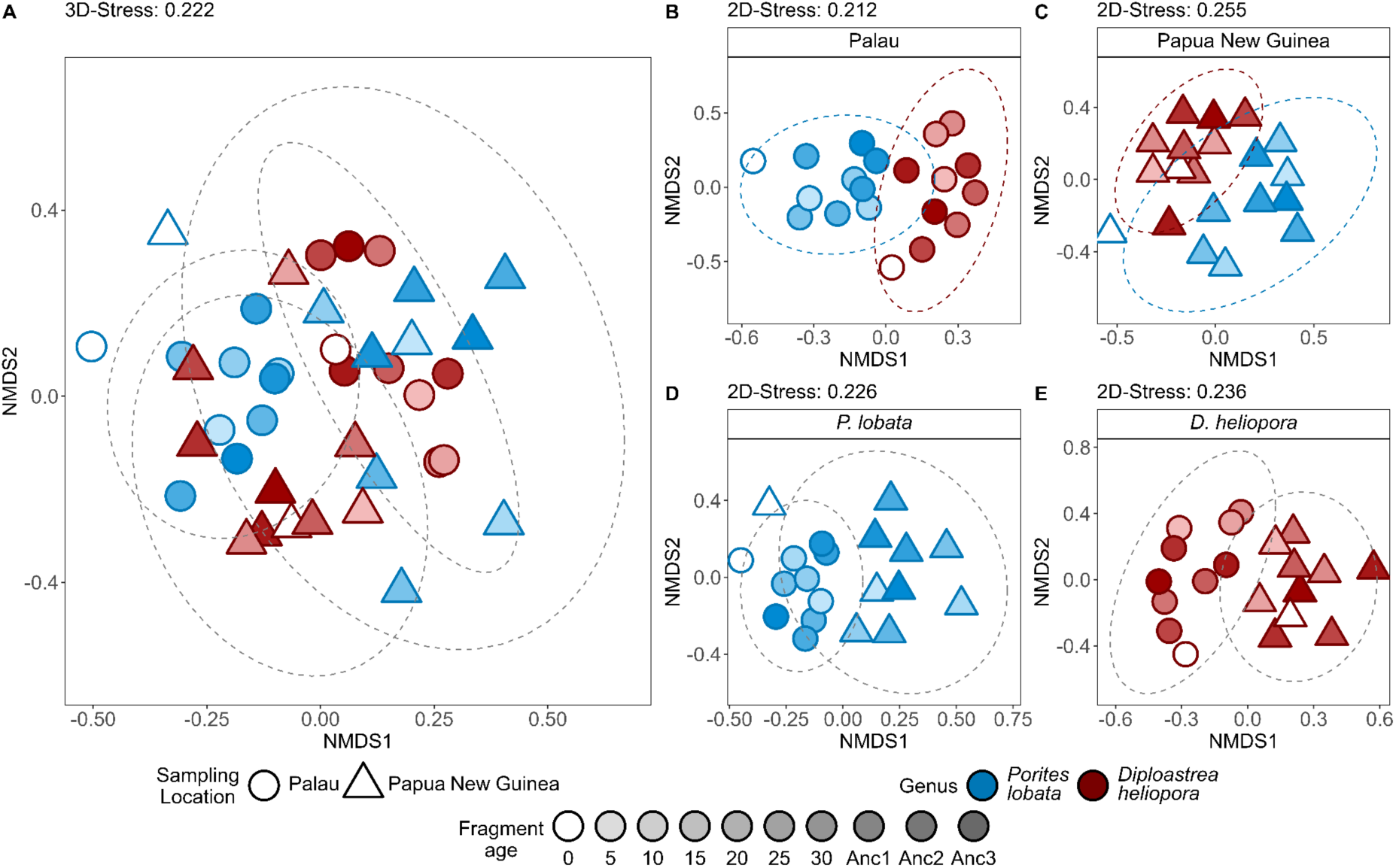
Symbiodiniaceae community composition based on non-metric Multidimensional Scaling of Bray-Curtis distances of post–MED sequences. Ellipses represent 95 % CI. Symbiodiniaceae communities **A.** from both coral species and sampling locations, **B.** from *P. lobata* and *D. heliopora* sampled in Palau, **C.** in Papua New Guinea, and **D.** from *P. lobata,* and **E.** *D. heliopora* at both locations.

Reconstructed Symbiodiniaceae communities from Papua New Guinea were more stable over decadal and centennial timeframes (Fig. 4A). With the exception of the most recent sample from the top of the cores, no shifts in dominance between Symbiodiniaceae genera were observed, though sequences within genera varied. In contrast, shifts in the dominant symbiont genus were recorded in the cores from Palau (Fig. 4A). While the skeletons of both coral species were largely dominated by sequences belonging to the genus *Symbiodinium*, the 10- year old sample from *P. lobata* was dominated by *Cladocopium* sequences. The 30-year old sample of *D. heliopora* was dominated by *Durusdinium* sequences and the sequences recovered from the 25-year old sample indicate a trend where *Durusdinium* dominance slowly reversed back to *Symbiodinium* dominance that had reestablished in the 20-year old sample (Fig. 4A). These site-specific patterns were also reflected in a higher number of shared post-MED sequences for the cores from Papua New Guinea (80) when compared to Palau (23) (Fig. S1).

**Figure. 4.**
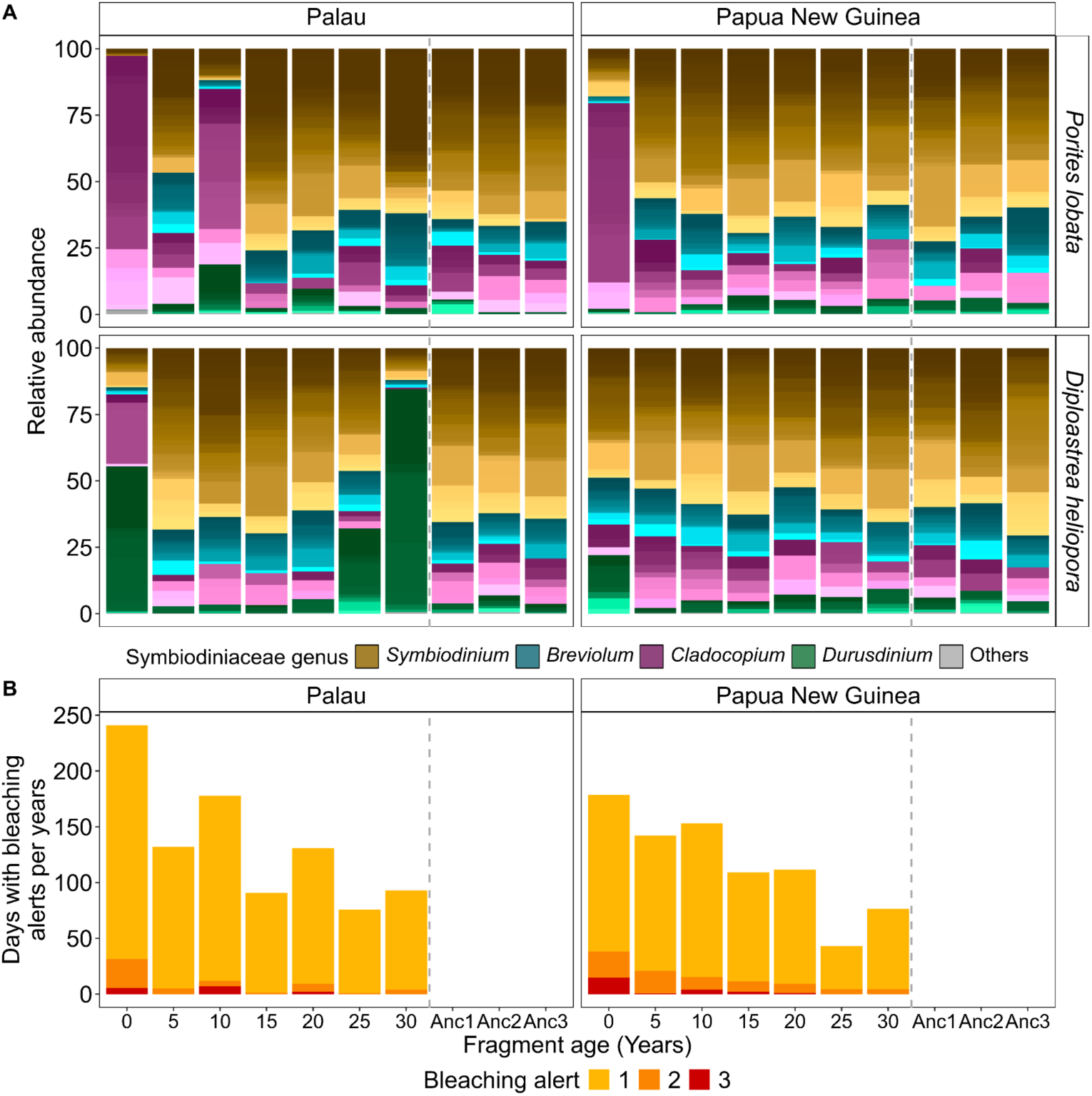
Symbiodiniaceae communities recovered from coral skeleton cores in relation to historical bleaching alerts. **A.** Relative abundance of Symbiodiniaceae genera of *P. lobata* and *D. heliopora* sampled in Palau and Papua New Guinea. Color shades represent unique post- MED sequences. **B.** Daily bleaching alert reconstructions from sea surface water temperatures for the time intervals corresponding with the coral core fragment ages.

### 3.3. Symbiodiniaceae community dynamics partially coincided with historical heat stress events

The historical SST data obtained from the NOAA Coral Reef Watch revealed increasing frequency and severity of SST anomalies associated with bleaching alerts in both locations over time, which peaked at over 220 and 150 bleaching alert days for the sampling year of the cores for Palau and Papua New Guinea, respectively (Fig. 4B, Fig. S2). In general, a higher number of annual bleaching alert days were recorded at Palau than at Papua New Guinea. The occurrence of a severe alert level 3 bleaching episode in Palau coincided with a change in dominant symbiont in *P. lobata* (Fig. 4), while the changes observed in *D. heliopora* could not be matched to a distinct event based on the NOAA bleaching alert data.

## 4. Discussion

### 4.1. aDNA extraction protocol and sample decalcification affect Symbiodiniaceae community reconstructions

Reconstruction of Symbiodiniaceae communities was successful at decadal and centennial-scale from *P. lobata* skeleton cores with all DNA extraction methods. The main differences between the tested protocols included the use of a decalcification step with EDTA prior to the extraction in two out of three methods and the use of a commercial extraction kit vs. a bespoke protocol for ancient DNA. The protocols that included the decalcifications step shared the dominance of *Cladocopium* sequences in the core top sample as well as the higher number of shared post-MED sequences. These results suggest that the dissolution of the aragonite (CaCO3) matrix yielded more comparable results regardless of the subsequent extraction protocol. The use of a decalcification step is common for aDNA extractions of coral skeletons, other marine calcifying invertebrates including mollusk shells, foraminifera, and mammal bones to release DNA bound to the skeletal architecture (Waller et al. 2007; Lendvay et al. 2020; Scott et al. 2022; González-Pech et al. 2024; Martin-Roy et al. 2024). Nevertheless, aDNA extractions employing coral skeletons have been successfully performed without the addition of a decalcification step, which is also supported by our data (Connelly et al. 2024; Rey et al. 2025). Another difference between protocols was the lower starting material in the Ancient DNA approach of 50 mg vs. 250 mg in the other protocols. However, these subsamples were all taken from the same powdered stock material, minimizing batch effects. In addition, despite differences in input mass, the two protocols that included a decalcification step yielded more similar Symbiodiniaceae communities than those of the two protocols with the same input mass. Together, these data highlight that a fraction of Symbiodiniaceae DNA is bound within the aragonite skeleton and that a decalcification step before DNA extraction enables the capture of this diversity regardless of which protocol is subsequently used for extraction.

Commercially available DNA extraction kits have previously been used for aDNA extraction of invertebrate aragonite skeletons. The PowerSoil Pro kit was selected for testing due to the inclusion of guanidine hydrochloride and silica spin columns that prevent the co-purification of PCR inhibitors common in environmental samples (Martin et al. 2021; Connelly et al. 2024; Martin-Roy et al. 2024). While lower DNA yields were reported with this kit when compared to the Blood and Tissue DNA kit (Qiagen, Germany) (Connelly et al. 2024), the combination with the decalcification step provided a higher amplification success in our early testing. This suggests that commercial kits may provide a simple alternative to the use of specialized aDNA extraction protocols with user-made buffers without a loss of reconstructed Symbiodiniaceae communities at century scales. This result is supported by the successful reconstruction of communities from 170-year-old coral skeleton core fragments employing the QIAamp DNA Micro Kit (Qiagen, Germany) (Rey et al. 2025). Moreover, the use of the Ancient DNA protocol did not allow for the recovery of sufficient quality Symbiodiniaceae sequences from millenia- old coral (Scott et al. 2022), suggesting that even optimized protocols for the recovery of shorter DNA fragments common in old and poorly preserved samples may not provide additional benefits for Symbiodiniaceae reconstructions.

Taken together, our results show that the DNA extraction protocol has an effect on the recovered Symbiodiniaceae communities, which has implications for the comparability across studies. As such, we recommend the use of one consistent DNA extraction method with a decalcification step for all samples of a study to capture the additional diversity. Moreover, additional DNA extraction optimizations that improved yields in forensic mammalian bones could be evaluated in further studies. These include testing DNA concentrations in the EDTA decalcification solution and DNA enzymatic repair techniques to test their effects in the reconstructed Symbiodiniaceae communities from coral skeletons (Hofreiter et al. 2021).

### 4.2. aDNA from coral cores recovers diverse Symbiodiniaceae communities

Different Symbiodiniaceae communities were reconstructed from all cores, highlighting the community specificity of each coral species and between geographic locations. Coral species are known to associate with specialized Symbiodiniaceae communities (Davies et al. 2023) and within species, these communities may vary along spatial, temporal, or environmental gradients (Terraneo et al. 2019; De Souza et al. 2022; Lewis et al. 2024; Ng et al. 2024). Taken together, our results indicate that the reconstructions are sensitive to changes associated with taxonomic and geographical factors, a novel finding that previous Symbiodiniaceae community reconstructions have not evaluated (González-Pech et al. 2024; Rey et al. 2025).

The recovered Symbiodiniaceae genera partially correspond to modern coral tissue samples. The dominance of *Cladocopium* in the tissue fragments of both *P. lobata* cores and the high relative abundance of *Durusdinium* in the tissue fragment of *D. heliopora* sampled in Palau is consistent with samples from other reefs of the Indo-Pacific including in Palau and Papua New Guinea during the Tara Pacific expedition as well as long-term community monitoring studies (Hume et al. 2020; Lewis et al. 2024; Ng et al. 2024). Nevertheless, *Symbiodinium* was found in high abundances (> 50 %) in most samples below the core top regardless of coral species and sampling location followed by a lower (∼10 %) but consistent abundance of *Breviolum*. The genus *Symbiodinium* has been detected to be present mostly as a low-abundant background and rarely as dominant symbiont, in multiple coral species including *P. lobata* and *P. lutea* sampled in Palau and Papua New Guinea (Smith et al. 2017; Hume et al. 2020; Ng et al. 2024), partially explaining its occurrence in the skeleton samples. Notably, the genus *Breviolum* is more frequently reported from Atlantic coral reefs, where it has also been reconstructed from coral cores of *Orbicella faveolata* (Garrido et al. 2022; González-Pech et al. 2024). In the Pacific, it has only occasionally been associated with corals, reef water, and sediments in the Great Barrier Reef, with other cnidarians such as hydrocorals and anemones in the eastern and central Pacific, and with the coral *Plesiastrea versipora* in the northern temperate Pacific (LaJeunesse et al. 2018; Ziegler et al. 2018, 2019; Nitschke et al. 2022). Similar to finding *Symbiodinium* in the coral skeletons, while not implausible, the detection of *Breviolum* raises questions about the process of Symbiodiniaceae deposition in coral skeletons.

There is currently no established model of Symbiodiniaceae DNA deposition in coral skeletons. The coral skeletal matrix is a low-light environment that experiences periodic anoxic conditions (Iha et al. 2021). Shallow coral skeletal bands may support a distinct endolithic community often composed of green algae from the genus *Ostreobium*, fungi, and bacteria (Tribollet et al. 2006; Iha et al. 2021). These endolithic organisms have adaptations to the challenging conditions within the skeleton, which Symbiodiniaceae cells do not seem to possess (Iha et al. 2021). One possible mechanism involves the accidental entrapment of the native Symbiodiniaceae communities from the coral tissue in the skeleton, e.g., when horizontal dissepiments are deposited during calcification or after injury. Whether and for how long these symbiont cells remain viable is unknown. A complementary explanation involves the entrapment of Symbiodiniaceae DNA in the aragonite matrix of the coral skeleton similar to the preferential DNA preservation in the columnar aragonite CaCO3 layer of mollusk shells (Martin et al. 2021). Additionally, mollusk shells capture and preserve environmental DNA (Der Sarkissian et al. 2017). The transport of seawater minerals to the calcifying fluid of corals might provide an explanation for the preservation of exogenous DNA in skeletons, although possible transport mechanisms need to be investigated (Tambutté et al. 2011; Der Sarkissian et al. 2017; Thompson 2022). Our results suggest the deposition of multiple Symbiodiniaceae genera in coral skeletons. Further studies are needed to elucidate the underlying mechanisms that lead to the mismatch in relative abundances between genera between coral tissue and the underlying skeleton in aDNA derived communities. For instance, the Symbiodiniaceae cells may use the skeleton as a habitat, or the cells or only their DNA may be trapped. An example of entrapment of exogenous organisms is the record of the dinoflagellate *Azadinium spinosum* in coral skeleton from aDNA-based reconstructions in *P. lobata*. *Azadinium spinosum* is not commonly associated with corals and it has been hypothesized that its traces were incorporated in the skeleton from the surrounding seawater by ingestion during algae bloom events (Rey et al. 2025). The presence of diverse Symbiodiniaceae communities in the seawater (Claar et al. 2020; Million et al. 2024) may thus explain their detection in coral skeletons and the record of these communities in coral skeletons highlight the potential to explore the use of coral skeletons for environmental DNA (eDNA) based reconstructions in time.

### 4.3. Reconstructions from coral cores capture past Symbiodiniaceae community dynamics

Symbiodiniaceae communities in cores from Papua New Guinea were relatively stable at decadal and centennial time scales and only the tissue-associated core tops were distinct from the older skeletal samples. In contrast, both cores sampled in Palau suggest changes in the dominant symbiont community during the last decades. Notably, the large shifts in symbiont community composition between time points were characterized by dynamics of sequences within the extant community and novel sequences, indicating a combination of symbiont shuffling and shifting as the underlying mechanism (Quigley et al. 2022).

Notably, overall higher SST and frequency of days with severe bleaching alerts were observed in Palau when compared to Papua New Guinea, suggesting a larger influence of historical heat stress events that could be associated with Symbiodiniaceae community dynamics (Lewis et al. 2024; Ng et al. 2024). The higher frequency of days with severe bleaching alerts coincided with the dominance of *Cladocopium* over *Symbiodinium* in *P. lobata* within the first 10 years of the reconstruction, which is similar to previously reported patterns in *O. faveolata* reconstructions (González-Pech et al. 2024). While *Porites* spp. are often tightly associated with the C15-type complex of *Cladocopium*, *P. lobata* was previously shown to undergo a lab-induced symbiont switch (Scharfenstein et al. 2022), as well as a high spatio-temporal structure in the reef (Ziegler et al. 2015), deeming a symbiont change plausible.

The occurrence of a severe alert level 3 bleaching episode in Palau coincided with a change in dominant symbiont in *P. lobata*. While the *Durusdinium* dominance in *D. heliopora* in Palau could not be linked to a NOAA bleaching alert, this sample was taken following the El Niño-associated bleaching in 1982/1983 (Coffroth et al. 1990). Given the low spatial and temporal resolution in environmental history and proof-of-concept sampling design, the causal link between environmental history and community dynamics remains to be confirmed. The subsequent sample from 5 years later indicates a lower but still dominant *Durusdinium* presence, which had reversed back to *Symbiodinium* dominance in the skeleton another 5 years later. This could reflect the reversal in symbiont communities to their original state, as reported from the field (Smith et al. 2017; Quigley et al. 2022). The coral skeleton-based Symbiodiniaceae community reconstruction cannot unequivocally confirm past changes of tissue-associated communities due to a lack of understanding of DNA entrapment mechanisms. However, as suggested by our data, reconstructions reveal patterns that may indicate changes in dominance of the community that were previously inaccessible due to a lack of long-term time-series samples. Moreover, by combining past ocean chemistry and climate reconstructions from the same cores employed for aDNA sampling, high resolution correlations between Symbiodiniaceae dynamics and past environmental conditions may be established. The information derived from such multidisciplinary studies may be used to better inform projections of coral health and status under future ocean conditions.

### 4.4. Perspective

Ancient DNA approaches to reconstruct microbial dynamics or host adaptation processes from coral skeletons are challenged by the environmental conditions during the lifetime of the coral that favor DNA degradation. In this study, aDNA as far back as 115 years was successfully recovered, validating the use of coral skeleton records for comparative analyses of long-term Symbiodiniaceae community dynamics. Similar to the reconstruction of past ocean conditions from geochemical proxies, the reconstruction of Symbiodiniaceae communities is limited by the available skeletal material, which often lacks biological replicates per species and site. Further research on the mechanisms of aDNA deposition in the coral skeleton will help to interpret reconstruction dynamics and to understand natural long-term adaptations to environmental stressors.

## Author contributions

JFG: performed research, analyzed data, wrote the paper, VT: performed research, JR: performed research, MC: performed research, SR: contributed reagents or analytical tools, ED: contributed reagents or analytical tools, MZ: designed research, contributed reagents or analytical tools, wrote the paper

## Acknowledgements

Special thanks to the Tara Ocean Foundation, the R/V Tara crew and the Tara Pacific Expedition Participants (https://doi.org/10.5281/zenodo.3777760). We are keen to thank the commitment of the following institutions for their financial and scientific support that made this unique Tara Pacific Expedition possible: CNRS, PSL, CSM, EPHE, Genoscope, CEA, Inserm, Université Côte d’Azur, ANR, agnès b., UNESCO-IOC, the Veolia Foundation, the Prince Albert II de Monaco Foundation, Région Bretagne, Billerudkorsnas, AmerisourceBergen Company, Lorient Agglomération, Oceans by Disney, L’Oréal, Biotherm, France Collectivités, Fonds Français pour l’Environnement Mondial (FFEM), Etienne Bourgois, and the Tara Ocean Foundation teams. Tara Pacific would not exist without the continuous support of the participating institutes. The authors also particularly thank Serge Planes, Denis Allemand, and the Tara Pacific consortium. The authors wish to thank Silvia Nachtigal (JLU) for support with lab work and the Marine Holobiomics Lab (JLU) for discussions of the data. This is publication number #40 of the Tara Pacific Consortium.

## Supplementary information

**Table S1.** Post-MED sequence number of each core fragment for *P. lobata* and *D. heliopora* sampled in Palau and Papua New Guinea. The counts represent the remaining post-MED sequences kept after blank correction.

| Sample | Fragment age | Post-MED sequence number | Sample | Fragment age | Post-MED sequence number |
| --- | --- | --- | --- | --- | --- |
| Palau -<br>*P. lobata* | 0 | 59784 | Papua New Guinea -<br>*P. lobata* | 0 | 35064 |
|  | 5 | 28479 |  | 5 | 24862 |
|  | 10 | 36229 |  | 10 | 36820 |
|  | 15 | 29205 |  | 15 | 33140 |
|  | 20 | 33067 |  | 20 | 37420 |
|  | 25 | 36417 |  | 25 | 54964 |
|  | 30 | 33104 |  | 30 | 34415 |
|  | 80 | 33654 |  | 80 | 24150 |
|  | 85 | 29402 |  | 85 | 35839 |
|  | 90 | 38003 |  | 90 | 29501 |
| Palau -<br>*D. heliophora* | 0 | 108490 | Papua New Guinea -<br>*D. heliophora* | 0 | 54359 |
|  | 5 | 22595 |  | 5 | 35561 |
|  | 10 | 30545 |  | 10 | 42582 |
|  | 15 | 22357 |  | 15 | 41390 |
|  | 20 | 21556 |  | 20 | 24405 |
|  | 25 | 55444 |  | 25 | 42712 |
|  | 30 | 110234 |  | 30 | 40984 |
|  | 100 | 31621 |  | 100 | 42819 |
|  | 105 | 37108 |  | 105 | 16940 |
|  | 110 | 44270 |  | 110 | 42416 |

**Table S2.** Pairwise statistical test of differences in the reconstructed Symbiodiniaceae communities obtained with three DNA extraction protocols from a *P. lobata* core sampled in Papua New Guinea. Pairwise PERMANOVA on Bray-Curtis distances with 999 permutations. DF: degrees of freedom, SS: sum of squares, F value: F statistic.

| Group comparison | Df | SS | F value | R<sup>2</sup> | Adj. p-value |
| --- | --- | --- | --- | --- | --- |
| PowerSoil vs.<br>Decalcification PowerSoil | 1 | 0.7899 | 1.8234 | 0.0920 | 0.003 |
| PowerSoil vs.<br>Decalcification Ancient DNA | 1 | 0.8508 | 2.0162 | 0.1007 | 0.006 |
| Decalcification PowerSoil vs.<br>Decalcification Ancient DNA | 1 | 0.6222 | 1.4443 | 0.0743 | 0.012 |

**Table S3.** Statistical test of multivariate dispersion in the reconstructed Symbiodiniaceae communities obtained with three DNA extraction protocols from a *P. lobata* core sampled in Papua New Guinea. Betadisper on Bray-Curtis distances. DF: degrees of freedom, SS: sum of squares, F value: F statistic.

| Factor | Df | SS | Mean Sq | F value | p-value |
| --- | --- | --- | --- | --- | --- |
| Groups | 2 | 0.001527 | 0.00076364 | 0.5064 | 0.6083 |
| Residual | 27 | 0.040717 | 0.00150805 |  |  |
| Total | 29 | 0.042244 |  |  |  |

**Table S4.** Statistical test of differences in the reconstructed Symbiodiniaceae communities of *P. lobata* and *D. heliopora* sampled in Palau and Papua New Guinea. PERMANOVA on Bray- Curtis distances with 999 permutations. DF: degrees of freedom, SS: sum of squares, Pseudo- F: pseudo-F statistic.

| Factor | Df | SS | R<sup>2</sup> | Pseudo-F | p-value |
| --- | --- | --- | --- | --- | --- |
| Coral species | 1 | 0.6966 | 0.0407 | 1.6932 | 0.0001 |
| Sampling location | 1 | 0.6720 | 0.0393 | 1.6334 | 0.0001 |
| Species x Location | 1 | 0.9176 | 0.0537 | 2.2304 | 0.0001 |
| Residual | 36 | 14.8110 | 0.8663 |  |  |
| Total | 39 | 17.0972 | 1.0000 |  |  |

**Table S5.**
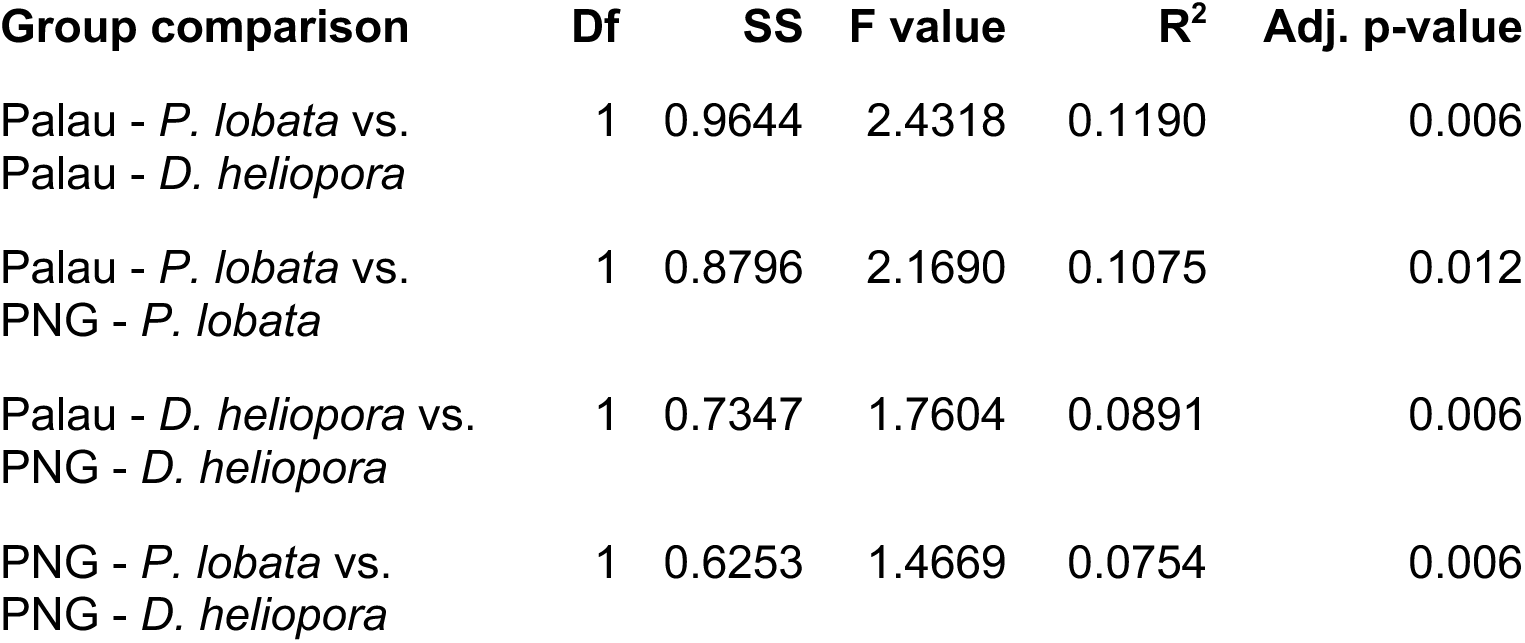
Pairwise statistical test of differences in the reconstructed Symbiodiniaceae communities of *P. lobata* and *D. heliopora* sampled in Palau and Papua New Guinea. Pairwise PERMANOVA on Bray-Curtis distances with 999 permutations. DF: degrees of freedom, SS: sum of squares, F value: F statistic.

| Group comparison | Df | SS | F value | R<sup>2</sup> | Adj. p-value |
| --- | --- | --- | --- | --- | --- |
| Palau - *P. lobata* vs.<br>Palau - *D. heliopora* | 1 | 0.9644 | 2.4318 | 0.1190 | 0.006 |
| Palau - *P. lobata* vs.<br>PNG - *P. lobata* | 1 | 0.8796 | 2.1690 | 0.1075 | 0.012 |
| Palau - *D. heliopora* vs.<br>PNG - *D. heliopora* | 1 | 0.7347 | 1.7604 | 0.0891 | 0.006 |
| PNG - *P. lobata* vs.<br>PNG - *D. heliopora* | 1 | 0.6253 | 1.4669 | 0.0754 | 0.006 |

**Table S6.** Statistical test of multivariate dispersion in the reconstructed Symbiodiniaceae communities of *P. lobata* and *D. heliopora* sampled in Palau and Papua New Guinea. Betadisper on Bray-Curtis distances. DF: degrees of freedom, SS: sum of squares, F value: F statistic.

| Factor | Df | SS | Mean Sq | F value | p-value |
| --- | --- | --- | --- | --- | --- |
| Groups | 3 | 0.0173 | 0.0058 | 3.9191 | 0.0161 |
| Residual | 36 | 0.0530 | 0.0015 |  |  |
| Total | 39 | 0.0703 |  |  |  |

**Figure S1.**
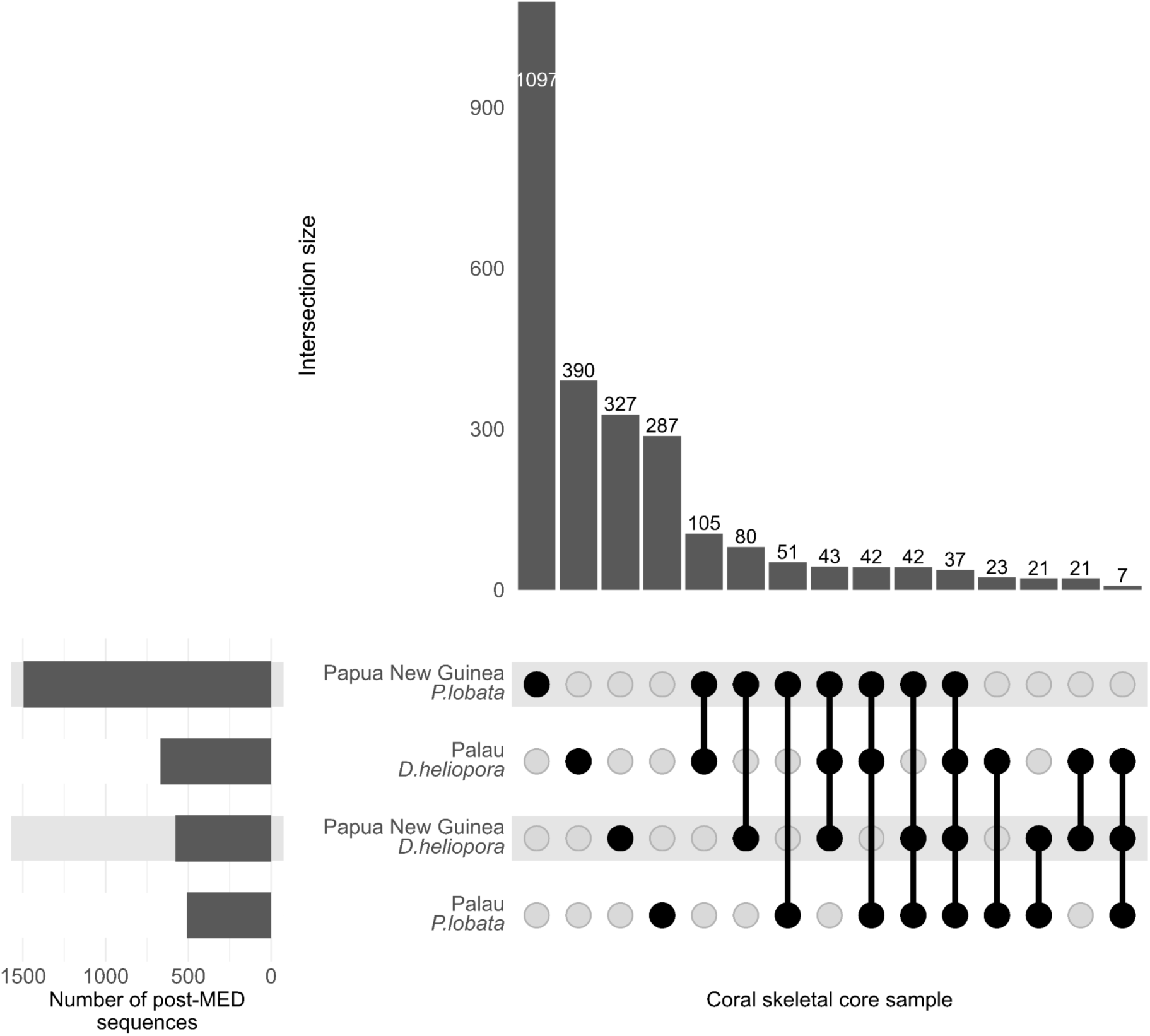
Number of shared post-MED sequences between *P. lobata* and *D. heliopora* skeletal cores sampled in Palau and Papua New Guinea. The intersection size (vertical bar plot) represents the number of shared sequences of a set of skeletal core samples represented by filled points.

**Figure S2.**
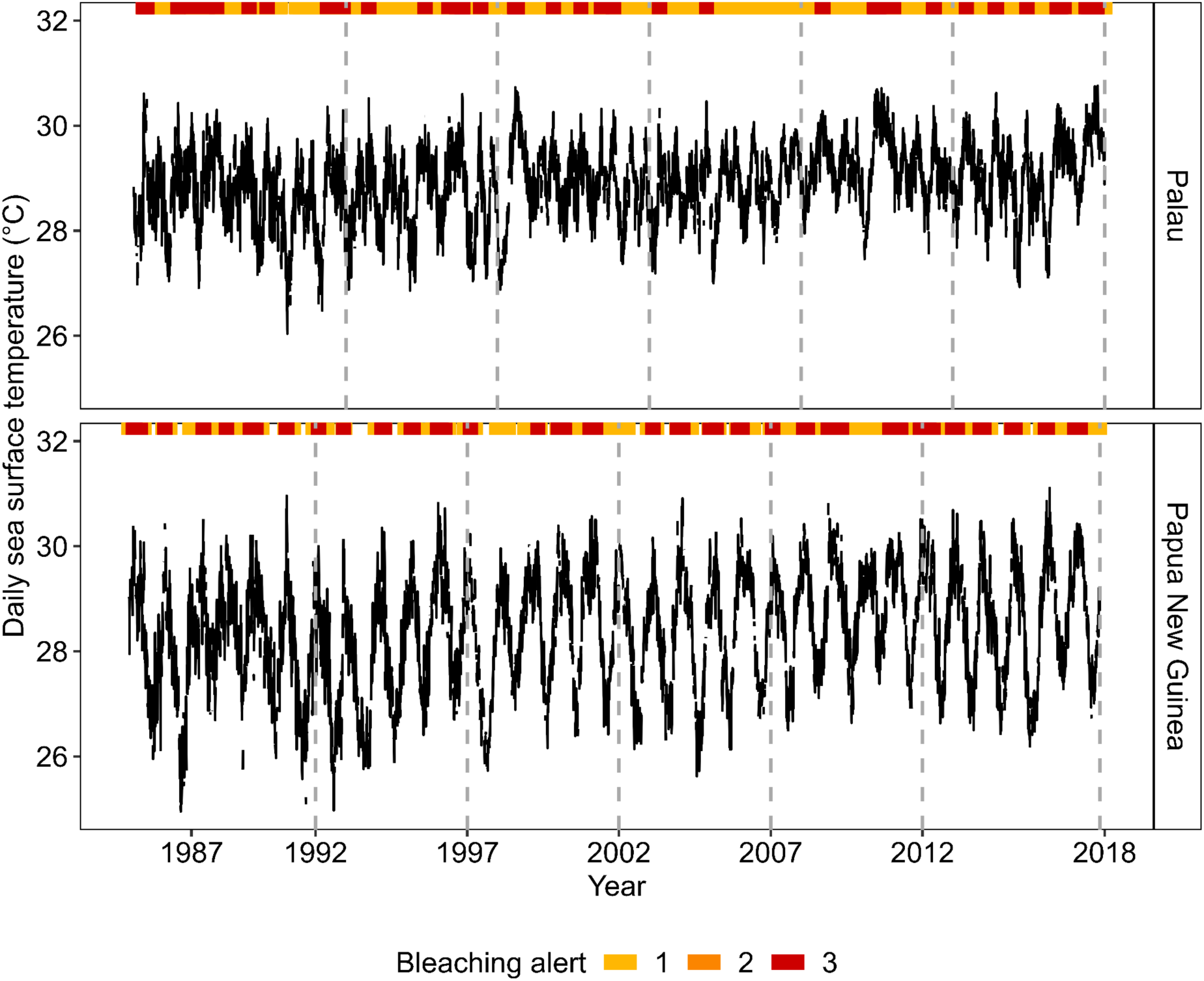
Daily sea surface temperature and days with bleaching alerts for the sampling locations obtained from the NOAA Coral Reef Watch database. The records range from 1985 to 2018.

